# A limbal stem cell deficiency murine model with residual limbal stem cells

**DOI:** 10.1101/2024.10.17.618942

**Authors:** Hideaki Someya, Shintaro Shirahama, Margarete M. Karg, Meredith S. Gregory-Ksander, Reza Dana, Bruce R. Ksander

## Abstract

Bilateral limbal stem cell deficiency (LSCD) is a significant cause of corneal blindness and is more difficult to treat, as compared with unilateral LSCD because no source of autologous limbal stem cells (LSCs) remains in these patients. Thus, bilateral patients could be candidates for treatment with allogeneic LSC transplants that require long-term systemic immunosuppression therapy. Thus, if possible, for the correct candidates, using autologous LSCs could be a preferred treatment. Recent *in vivo* laser confocal microscopic examination of the ocular surface *in situ*, combined with impression cytology, has indicated that some patients diagnosed with a complete bilateral LSCD possess residual LSCs. However, it remains unknown whether these residual LSCs still have stem cell potential due to the lack of animal models that mimic this pathology. The goal of the current study is to make a complete LSCD model that possesses evidence of residual LSCs. We induced complete LSCD in mice using two methods: (1) removed the corneal epithelium and the epithelial basement membrane using a rotating burr, and (2) removed the corneal epithelium using 20% ethanol but retained an intact epithelial basement membrane. A complete LSCD was defined by a lack of CK12-positive corneal epithelial cells and the presence of infiltrating CK19-positive conjunctival epithelial cells. Corneas were examined for wound closure, corneal opacity, LSC exhaustion, and inflammation. We observed that complete LSCD mice without an intact epithelial basement membrane resulted in few residual LSCs. By contrast, complete LSCD mice that retained the epithelial basement membrane were accompanied by a reduced inflammatory response plus a significant number of residual LSCs. This model will allow future studies to determine the function of residual LSCs in complete LSCD.

## 1. Introduction

There are a variety of different causes of limbal stem cell deficiency (LSCD) (reviewed in Deng et al. 2019), which can be either partial or complete, unilateral or bilateral. Ophthalmologists have made significant advances in the diagnosis and management of LSCD patients, and the International LSCD Working Group published new guidelines on the definition, classification, diagnosis, staging, and management of LSCD (Deng et al. 2019, 2020). The consensus is that, due to the extensive heterogeneity of LSCD patients, their treatment must be geared to the specific characteristics of the patient based on the cause, extent, severity, and stage of LSCD.

There are multiple treatment options for patients with unilateral LSCD that utilize limbal tissue from the normal contralateral eye (reviewed in Deng et al. 2020). These approaches have reported varying levels of success in treating unilateral LSCD. By contrast, patients with bilateral LSCD often have little residual autologous limbal stem cells (LSCs), and therefore investigators have treated these patients with transplants using allogeneic limbal tissue obtained from cadaveric donors (Deng et al. 2020; Satake et al. 2013), using either allogeneic whole tissue transplants or *in vitro* expanded allogeneic limbal tissue-derived cells. In general, however, these allogeneic transplants have a worse outcome than autologous transplants. These patients must receive long-term systemic immunosuppressive therapy, and failure of the transplants is possibly due to allograft immune rejection. For this reason, using autologous LSCs will always be the preferred treatment for LSCD patients.

One of the most important advances clinically in the diagnosis of LSCD is the use of *in vivo* laser confocal microscopy to examine the patients’ ocular surface *in situ*, which, combined with impression cytology, provides the most detailed information on the status of the corneal epithelium and limbus in these patients. Using this technique, Sophie Deng and coworkers identified areas of normal limbal epithelial cells in patients who, by clinical examination, were diagnosed with a complete bilateral LSCD (Chan et al. 2016). Moreover, in our own study, we observed that sections of limbus obtained from patients with a complete LSCD due to an alkali burn displayed ABCB5+ (limbal stem cell marker) cells by immunohistochemical staining (Ksander et al. 2014), indicating that LSCs could be found in the limbus of some patients with a complete LSCD. Clearly, whatever LSCs remain in these patients, they are unable to sustain any extensive production of corneal epithelial cells. Whether these residual LSCs are too few in number and/or functionally deficient is unknown. The goal of the current study is to induce a complete LSCD that possesses residual LSCs, which would allow future studies to determine their functional activity and explore methods to restore dysfunctional stem cell activity.

The most common approach for inducing an LSCD is a mechanical total limbus-to-limbus corneal de-epithelization using either a blunt blade or a rotating burr (Delic et al., 2022). For de-epithelization using a blunt blade, difficulty in scraping off the epithelium can lead to incomplete de-epithelization (Afsharkhamseh et al., 2016), resulting in a highly variable LSCD phenotype. For de-epithelization using a rotating burr, the rotating burr scrapes off the epithelial cells as well as, the epithelial basement membrane (EBM) (Li at al., 2016), leading to a more consistent and pronounced LSCD phenotype.

The application of ethanol to the corneal surface can detach the corneal epithelium at the level of the hemidesmosomes located between the epithelial basal cells and the EBM (Browning et al., 2003; Espana et al., 2003). In the clinical setting, the corneal epithelium is peeled using an eye spear after a circular sponge soaked in 20% ethanol is placed on the cornea (Dua et al., 2021). This method allows for scraping off the corneal epithelium more easily and leaving the EBM intact. In the current study, we used two techniques to induce a complete LSCD in mice: (i) mechanical de-epithelialization, where the EBM was removed, or (ii) ethanol de-epithelialization, where the EBM remained intact. Our results showed that removing the corneal epithelium with retained EBM resulted in a complete LSCD defined by a lack of CK12+ corneal epithelial cells and the presence of infiltrating CK19+ conjunctival epithelial cells. However, this occurred with a reduced inflammatory response and a significant number of residual LSCs. This model will allow future studies to determine the function of residual LSCs in complete LSCD.

## 2. Materials and methods

### 2.1. Mice

C57BL/6J mice (Strain #:000664) were obtained from The Jackson Laboratory (Bar Harbor, ME, USA). Mice were bred under normal circadian rhythms (a 12-h light/dark cycle) and allowed free access to food and water. All mice used for experiments were 8-10 weeks old. For anesthesia, ketamine (5 mg/ml) and xylazine (2 mg/ml) mixture solution was injected intraperitoneally for survival procedures (10μL/g body weight). For non-survival procedures, mice were euthanized using carbon dioxide inhalation followed by cervical dislocation. All murine experiments were performed in accordance with the statement for the use of animals in ophthalmic and vision research by the Association for Research in Vision and Ophthalmology. This study was approved by the Animal Care Committee of the Massachusetts Eye and Ear Infirmary (Protocol Number: 2022N000027).

### 2.2. Limbal stem cell deficiency model

To make an LSCD model that retained a complete EBM, we performed total corneal de-epithelization using ethanol. Before starting the procedure, we prepared circular cellulose filter paper (diameter: 1.5mm) and cylindrical cellulose filter paper (inner and outer diameter: 3 and 4 mm). First, after local anesthesia, we placed the circular filter paper and cylindrical filter paper on the cornea (Fig. 1A, B). Second, we kept dripping 20% ethanol (3.5μL) on the circular filter paper for thirty seconds (Fig. 1C). Third, after discarding the filter paper, we flushed the ocular surface twice with phosphate-buffered saline (PBS) to remove residual ethanol. Fourth, we started peeling off the central cornea epithelium using an eye spear (Fig. 1D, E), then extended the edges of the peeled area toward the limbus (Figs. 1F, G). Finally, to confirm the complete removal of the corneal epithelium, we stained the cornea with fluorescein and took corneal photographs using a cobalt blue filter using a DC-4 digital camera (Topcon, Tokyo, Japan) attached to an SL-D4 slit-lamp microscope (Topcon, Tokyo, Japan).

**Fig. 1.**
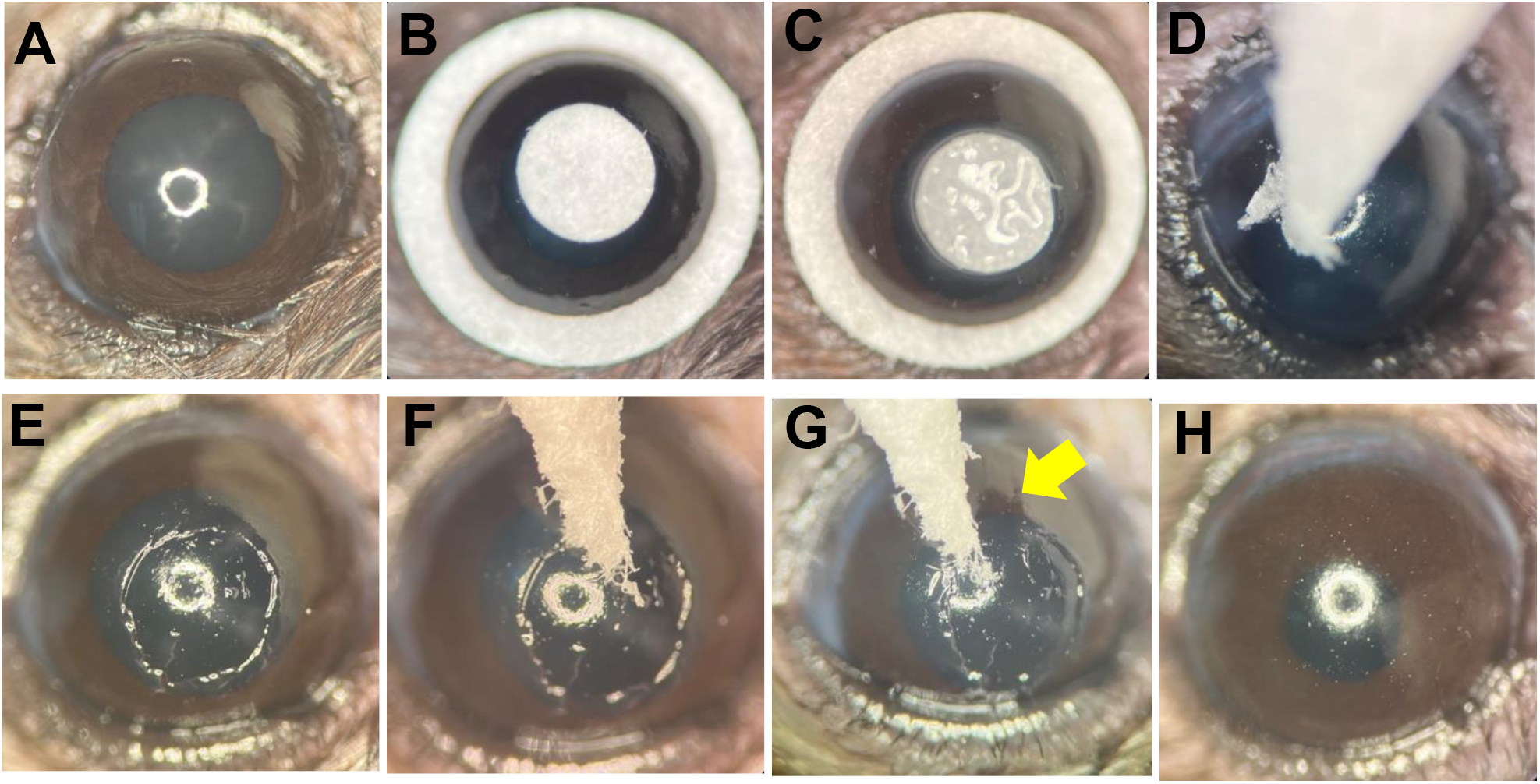
Total corneal de-epithelization using ethanol. (A) Before de-epithelization. (B) Place the circular filter paper (diameter: 1.5mm) on the central cornea and the cylindrical filter paper (inner and outer diameter: 3 and 4 mm) on the outside of the limbus. (C) Keep dripping the 20% ethanol (3.5μL) on the circular filter paper for thirty seconds. (D) Start peeling off the central cornea epithelium using an eye spear. (E) Completion of the central cornea de-epithelization. (F) Start extending the edges of the peeled area toward the limbus. (G) In the process of expanding the de-epithelization area. The yellow arrow indicates the boundary between peeled and unpeeled areas. (H) Completion of the total corneal de-epithelization.

To make a LSCD model that lacked an EBM, we performed total corneal de-epithelization using a rotating burr (Algerbrush II; Alger Inc., Lago Vista, TX, USA). After local anesthesia, we shaved the limbal epithelium circumferentially with a speed of 5 seconds per round. Then, we also shaved the epithelium from the limbus to the center in concentric circles using the rotating burr.

### 2.3. Central corneal de-epithelization

First, after local anesthesia, we placed the circular filter (diameter: 1.5mm) soaked with 20% ethanol on the central cornea for thirty seconds. Second, after discarding the filter paper, we flushed the ocular surface twice with PBS to remove residual ethanol. Third, we peeled off the central cornea epithelium using an eye spear. Finally, we stained the cornea with fluorescein to confirm the complete removal of the central corneal epithelium.

### 2.4. Evaluation of epithelial wound area and corneal opacity

We stained the cornea with fluorescein to evaluate the epithelial wound area. Then we took cornea photographs using a cobalt blue filter using DC-4 digital camera (Topcon, Tokyo, Japan) attached to an SL-D4 slit-lamp microscope (Topcon, Tokyo, Japan). The percentage of the fluorescein-stained area in the total cornea was evaluated by analyzing images using IMAGEnet6 software (Topcon, Tokyo, Japan). Based on the bright-field image, we evaluated the corneal opacity using a grading score from 0 to 4 (0 = completely clear, 4 = completely opaque) as previously described (Sonoda and Streilein, 1992).

### 2.5. Central corneal thickness measurement

Spectral-domain optical coherence tomography (SD-OCT) image was obtained using the Bioptigen Envisu R2200 Spectral Domain Ophthalmic Imaging System (Leica Microsystems, Wetzlar, Germany). Based on the acquired image, central corneal thickness (CCT) was measured using inVivoVue Diver 2.0 software (Leica Microsystems, Wetzlar, Germany).

### 2.6. Hematoxylin and eosin staining of corneal sections

Enucleated eyes were fixed in a 4% paraformaldehyde solution (Electron Microscopy Sciences, Hatfield, PA, USA) overnight and embedded in paraffin before thinly sliced (5μm). The sample was deparaffinized and stained with hematoxylin and eosin (H&E). Stained sections were photographed using a ZEISS Axio Imager M2 microscope

(Zeiss, Oberkochen, Germany).

### 2.7. Detection of slow-cycling cells

After performing total corneal de-epithelization, mice received an intraperitoneal injection (100μg/g body weight) of bromodeoxyuridine (BrdU) solution (Abcam, Cambridge, U.K.) (10mg/ml) for seven days. Eyes were collected 14 days after performing de-epithelization and immunostained by the method described below.

### 2.8. Immunostaining of cornea section

Eyes were enucleated and immediately embedded in Tissue-Tek^®^ O.C.T. Compound (SAKURA, Tokyo, Japan) and kept at -80°C until used. 15μm frozen sections were made using a Leica CM1950 cryostat (Leica Biosystems, Wetzlar, Germany). For immunostaining to detect laminin alpha 1/beta 1, cytokeratin 12 (CK12), and cytokeratin 19 (CK19), sections were fixed in 4% paraformaldehyde (Electron Microscopy Sciences, Hatfield, PA, USA) for 15 minutes at room temperature and then blocked in a blocking buffer (0.3% Triton X-100 (Sigma-Aldrich, St. Louis, MO, USA), 0.2% bovine serum albumin (BSA) (Sigma-Aldrich, St. Louis, MO, USA), and 5% goat serum (Sigma-Aldrich, St. Louis, MO, USA)) for 1 hour at room temperature and then incubated with primary antibody overnight at 4°C. After washing with 1×PBS, sections were incubated with the secondary antibody and 4′,6-diamidino-2-phenylindole (DAPI) solution (20μg/ml) (Thermo Fisher Scientific, Waltham, MA, USA) for 1 hour at room temperature. Finally, after washing with 1×PBS, sections were mounted using ProLong Diamond Antifade Mountant (Thermo Fisher Scientific, Waltham, MA, USA). Rabbit anti-keratin 12 monoclonal antibody (mAb) (clone: ERP17882) (1:100) (Abcam, Cambridge, U.K.), rat anti-cytokeratin 19 mAb (clone: TROMA-3) (1:200) (Sigma-

Aldrich, St. Louis, MO, USA), and rat anti-laminin alpha 1 / beta 1 mAb (1:100) (R&D Systems, Minneapolis, MN, USA) was used for the primary antibody. Alexa Flour 647-conjugated goat anti-rat IgG (H+L) antibody (1:2000) (Thermo Fisher Scientific, Waltham, MA, USA) and Alexa Flour 488-conjugated goat anti-rabbit IgG (H+L) antibody (1:2000) (Thermo Fisher Scientific, Waltham, MA, USA) was used for the secondary antibody.

For immunostaining to detect p63 and BrdU, sections were fixed in 4% paraformaldehyde solution for 15 minutes at room temperature and then blocked in a blocking buffer (0.3% Triton X-100, 0.2% BSA, and 5% goat serum) for 1 hour at room temperature and then incubated with rabbit anti-p63 mAb (clone: ERP5701) (1:200) (Abcam, Cambridge, U.K.) for 1 hour at room temperature. After washing with 1×PBS, sections were incubated with Alexa Flour 488-conjugated goat anti-rabbit IgG (H+L) antibody (1:2000) (Thermo Fisher Scientific, Waltham, MA, USA) for 1 hour at room temperature. After washing with 1×PBS, a second fixation was performed with 4% paraformaldehyde solution for 15 minutes at room temperature. Then, sections were soaked in 2N HCl solutions (Sigma-Aldrich, St. Louis, MO, USA) for 15 minutes at 40°C followed by a neutralization step with 1M borate buffer (pH=8.50) (Sigma-Aldrich, St.

Louis, MO, USA) for 10 minutes at room temperature. After washing with 1×PBS, sections were blocked in blocking buffer (0.3% Triton X-100, 0.2% BSA, and 5% goat serum) for 1 hour at room temperature and then incubated with rat anti-BrdU mAb (clone: BU1/75(ICR1)) (1:100) (Abcam, Cambridge, U.K.) overnight at 4°C. After washing with 1×PBS, sections were incubated with Alexa Flour 647-conjugated goat anti-rat IgG (H+L) antibody (1:2000) (Thermo Fisher Scientific, Waltham, MA, USA) and DAPI solution (20μg/ml) for 1 hour at room temperature. Finally, after washing with 1×PBS, all sections were mounted using ProLong Diamond Antifade Mountant.

### 2.9. RNA extraction and reverse transcription-quantitative polymerase chain reaction

Excised corneas were homogenized in a 1.5 ml tube with a pestle motor. After removing the debris, total RNA was extracted using RNeasy Plus Mini kit (Qiagen, Hilden, Germany). Total RNA quality was assessed using the absorbance ratio at 260 nm and 280 nm measured by NanoDrop 2000 Spectrophotometers (Thermo Fisher Scientific, Waltham, MA, USA). RNA samples with a 260/280 ratio of ≥ 2.0 were used in this study.

Total RNA was reverse transcribed into cDNA using SuperScript IV VILO Master Mix (Thermo Fisher Scientific, Waltham, MA, USA). The cDNA was amplified using both the specific primer sets listed in Table. S1 and Power SYBR Green PCR Master Mix (Thermo Fisher Scientific, Waltham, MA, USA) following the manufacturer’s instructions. The qPCR (cycling condition: 95°C 10 min and 40 cycles of [95°C 15 sec; 60°C 1 min]) was then performed using a CFX 384 Touch Real-Time PCR detection system (Bio-Rad, Hercules, CA, USA). Relative quantification was performed using the calibration curve method, and β-actin mRNA was used for transcript normalization. The relative RNA amount of each gene transcript in the cornea from EBM+ and EBM-groups was calculated based on that from the naive group.

### 2.10. Immunostaining image processing and analysis

Image capture was performed using Leica SP8 laser confocal microscope (Leica Microsystems, Wetzlar, Germany), and the analysis was performed using Image J software (National Institutes of Health, Bethesda, MD, USA). The % area for each marker was quantified as the percentage of each marker-stained area in the cornea epithelium region based on the DAPI image. To measure the p63/BrdU % area, we first extracted the co-localized area of p63 and BrdU. Then, the p63/BrdU % area was quantified as the percentage of the co-localized area in the cornea epithelium region determined in the same manner mentioned above. For % area calculation, we used the average value of the data acquired from the three different sections as the value for the individual sample.

### 2.11. Statistical analysis

Data were presented as mean ± standard deviation (SD) or standard error of the mean (SEM). The difference between the two groups was analyzed using Student’s t-test. Multiple-group comparisons wer analyzed using a one-way analysis of variance followed by the Tukey-Kramer test (\**p*<0.05, \*\**p*<0.01, \*\*\*\**p*<0.001, and \*\*\*\**p*<0.0001 throughout the paper). All statistical analyses were performed using JMP Pro 14 (SAS Institute Inc., Cary, NC, USA).

## 3. Results

### 3.1. Total corneal de-epithelization using ethanol retains the epithelial basement membrane

To assess complete corneal de-epithelization from limbus-to-limbus, using either a rotating burr or ethanol treatment, we stained the cornea with fluorescein and then took corneal photographs using a slit-lamp microscope. Both groups showed complete de-epithelization (Fig. 2A). Furthermore, to examine the depth of the corneal wound after de-epithelization, we measured the central corneal thickness (CCT) using an anterior segment OCT. CCT in both the ethanol and rotating burr groups was approximately 18μm thinner than in the naïve group (*p* < 0.001) (Fig. 2B, C). Finally, to assess the presence or absence of EBM after de-epithelization, we stained corneal sections using two methods (H&E staining and laminin alpha 1/ beta 1 immunostaining) and examined both the central cornea plus the limbal area. In both stained images, the ethanol-treated group retained a complete EBM (EBM+ group), while the rotating burr method group showed no evidence of the EBM (EBM-group) (Fig. 2D, E).

**Fig. 2.**
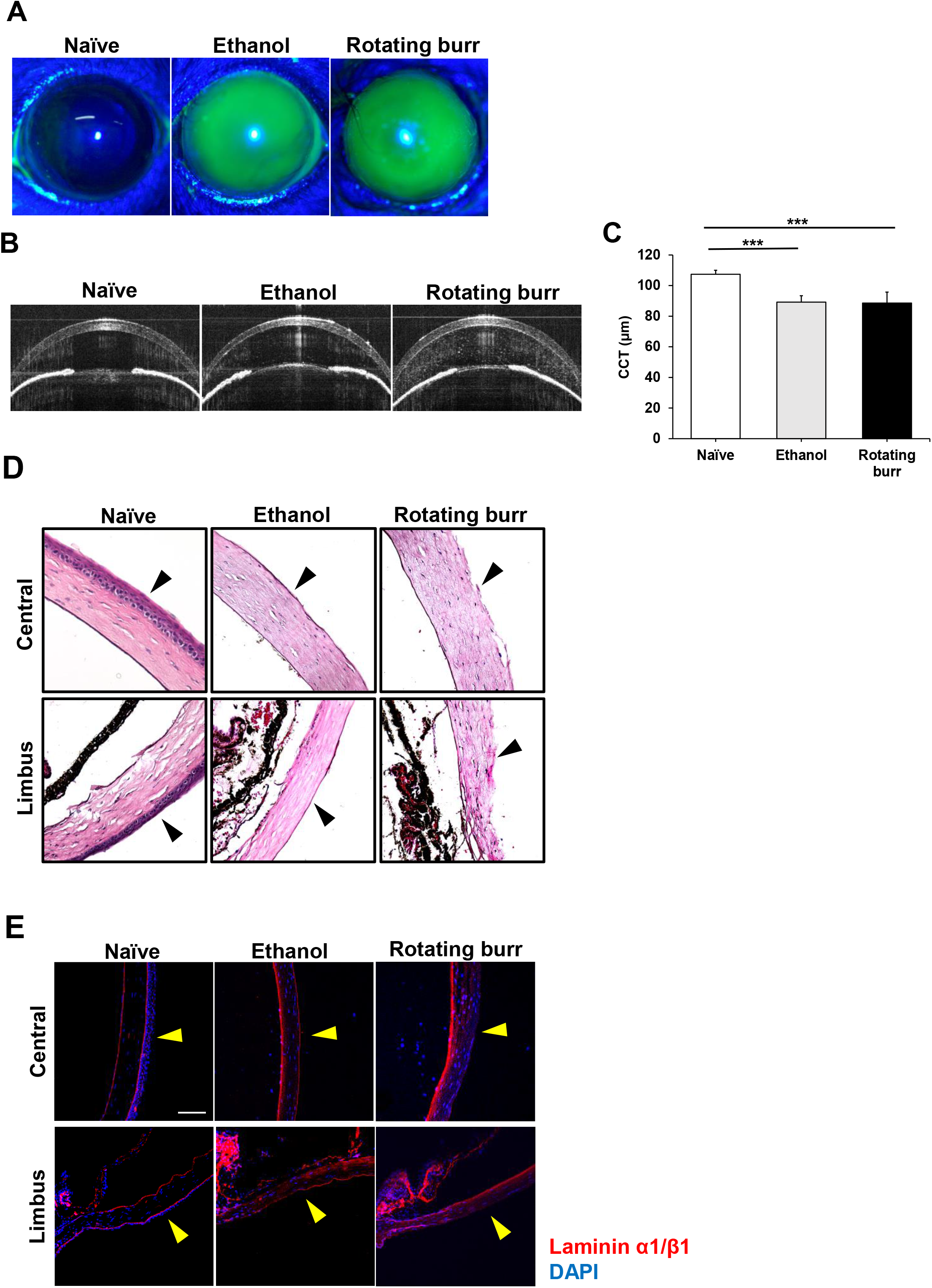
Total corneal de-epithelization using ethanol does not disrupt the epithelial basement membrane. (A) Representative fluorescein-stained cornea photographs immediately after total corneal de-epithelization by either ethanol treatment or using a rotating-burr. (B, C) Representative optical coherence tomography images showing the central cornea thickness (CCT) immediately after total corneal de-epithelization in ethanol treated or rotating-burr groups (n = 5 mice/group). Data are presented as mean ± S.D. Statistical analysis was performed using a one-way analysis of variance followed by the Tukey-Kramer test. \*\*\**p* < 0.001. (D) Representative hematoxylin and eosin-stained images immediately after total corneal de-epithelization in ethanol treated or rotating-burr groups. (E) Representative laminin alpha 1/beta 1 immunostaining images immediately after total corneal de-epithelization in ethanol treated or rotating-burr groups. The arrow indicates the epithelial side. Scale bars equal 100 μm.

### 3.2. The EBM promotes epithelial wound closure after total corneal de-epithelization

To evaluate the progress of wound closure and cornea opacity after total corneal de-epithelization, we took cornea photographs after de-epithelization every day for the first eight days and then on day 10, 12, and 14. For wound closure, the EBM+ group showed significantly faster wound closure (Fig. 3A, B). Specifically, the wounds closed by day 5 in the EBM+ group. In comparison, the wounds closed by day 10 in the EBM-group (Fig. 3A). We detected corneal opacity in both EBM+ and EBM-groups for seven days after de-epithelization (Fig. 3C, D). These results suggest that the presence of an EBM after de-epithelization is an important factor in promoting wound closure.

**Fig. 3.**
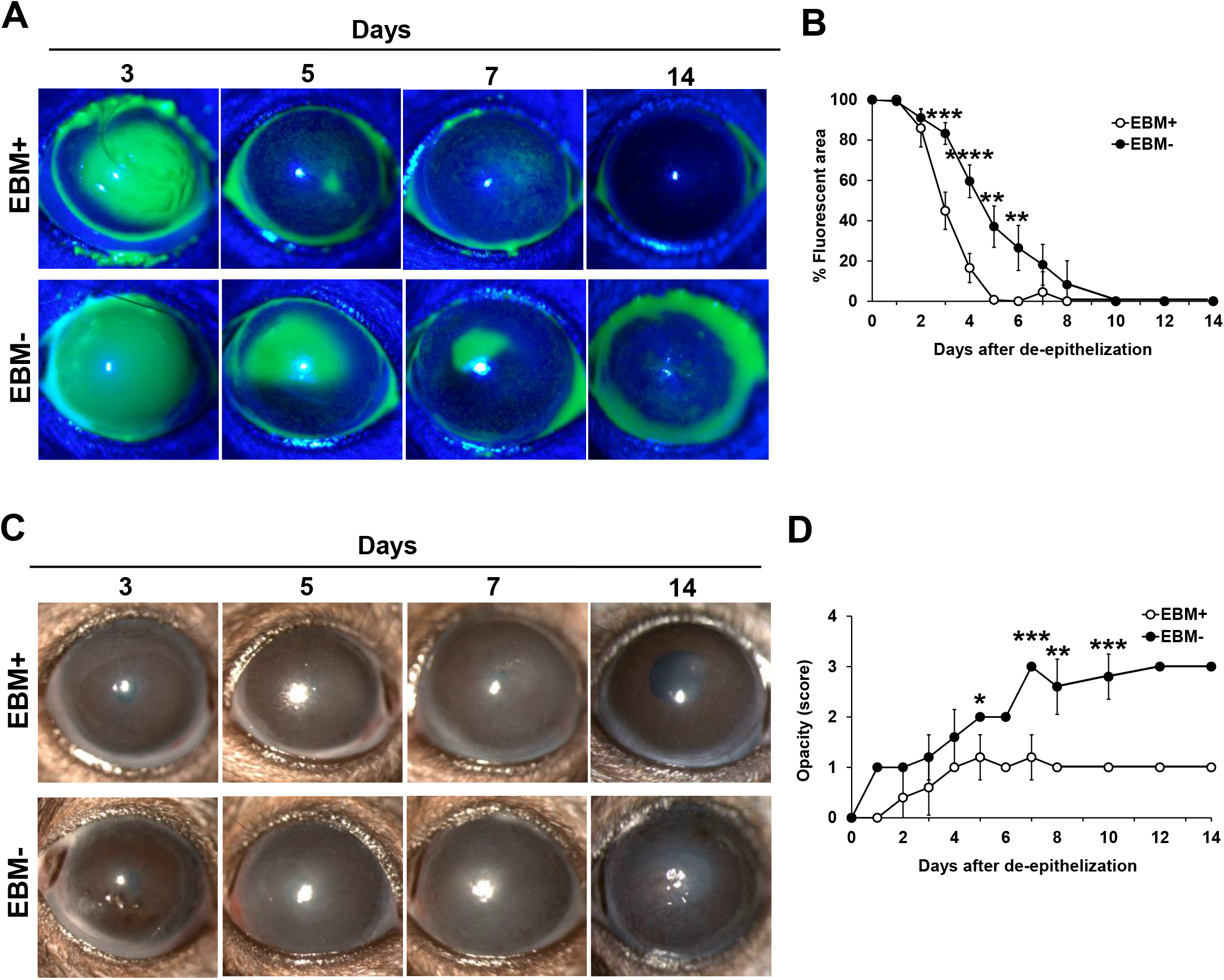
The presence of EBM after total corneal de-epithelization promotes corneal epithelial wound closure. (A, B) Representative fluorescein-stained cornea photographs on 3, 5, 7, and 14 days after total corneal de-epithelization in the EBM preserved (EBM+) and non-preserved (EBM-) groups. Quantitative graphs of the % fluorescein-stained area over time in the EBM+ and EBM-groups (n = 5 mice/group). (C, D) Representative cornea photographs on 3, 5, 7, and 14 days after total corneal de-epithelization in the EBM+ and EBM-groups. Quantitative graphs of the corneal opacity score over time in the EBM+ and EBM-groups (n = 5 mice/group). All data are presented as mean ± S.D. All statistical analyses were performed using Student’s t-test. \**p* < 0.05; \*\**p* < 0.01; \*\*\**p* < 0.001; \*\*\*\**p* < 0.0001. EBM, epithelial basement membrane.

In the experiment above, the wound closure observed following an LSCD was mediated by the infiltration of cells from the conjunctival epithelium. The following experiment was performed to determine whether the presence of an EBM facilitated wound closure by corneal epithelial cells when the limbus was intact. Mice received only a central corneal epithelial debridement by either ethanol treatment or rotating burr. As observed in mice with a complete LSCD, the central epithelial wound closed faster in the presence of an intact EBM as compared to the wound closure observed in mice without an EBM (Fig. S1A, B).

### 3.3. Total corneal de-epithelization with an intact EBM resulted in residual LSCs

LSCs are slow-cycling cells that rarely undergo cell division, resulting in the retention of labeled modified nucleic acids such as BrdU over a long period of time (Sartaj et al., 2017). Furthermore, p63 is also known as an epithelial stem cell marker (Pellegrini et al., 2001). Therefore, in this study, we identified LSCs as BrdU+ and p63+ co-localized cells (Lasagni et al., 2022). We performed total corneal de-epithelization using either ethanol treatment or a rotating burr, and then we collected the eyes 14 days later. To trace the LSCs, we injected the BrdU solution intraperitoneally for seven consecutive days after performing de-epithelization (Fig. 4A).

**Fig. 4.**
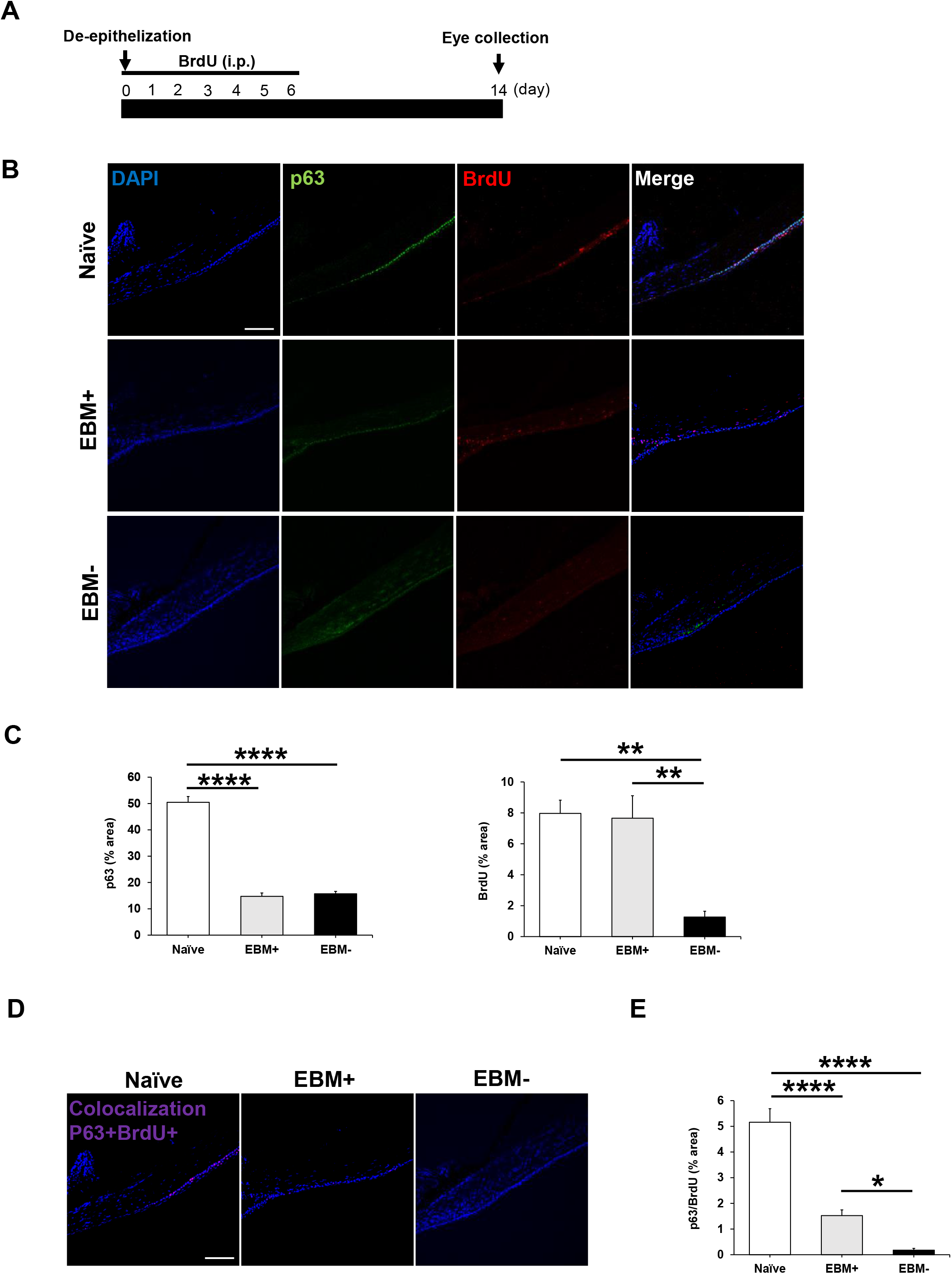
Total corneal de-epithelization with an intact EBM contains residual LSCs. (A) Schematic time course to detect limbal stem cells in the limbal stem cell deficiency murine model. (B, C) Representative p63 and BrdU immunostaining images 14 days after total corneal de-epithelization in the EBM+ and EBM-groups. Images show the limbus and peripheral cornea. Quantitative graphs of the % area for p63 and BrdU stained cells in this region in the EBM+ and EBM-groups (n = 5 mice/group). Scale bars equal 100 μm. (D, E) Representative images of the limbus and peripheral cornea where p63 and BrdU double-positive cells in the EBM+ and EBM-groups. Quantitative graphs of the % area for p63+ / BrdU+ co-localized cells in in the EBM+ and EBM-groups (n = 5 mice/group). Scale bars equal 100 μm. All data are presented as mean ± s.e.m. All statistical analyses were performed using a one-way analysis of variance followed by the Tukey-Kramer test. \**p* < 0.05; \*\**p* < 0.01; \*\*\*\**p* < 0.0001. EBM, epithelial basement membrane.

We evaluated the limbal and peripheral corneal epithelium for p63+ cells in EBM+ and EBM-groups, using naïve mice as a positive control (Fig. 4B, C). As expected, following induction of a LSCD in either the EBM positive or negative groups, there was a significant reduction in p63+ cells, as compared with naïve mice. Moreover, there was no significant difference between the reduction of p63+ cells between the EBM+ and EBM-groups (EBM+: 14.7 ± 1.3%, EBM-: 15.7 ± 0.9%) (Fig. 4B, C). By contrast, following induction of a LSCD that retained an intact EBM, there was no significant reduction in BrdU+ cells as compared to naïve mice (naïve: 8.0 ± 0.9%, EBM+: 7.7 ± 1.5%). In comparison, the induction of an LSCD without an EBM resulted in a significant loss of BrdU+ cells (84%) compared to naïve control mice. A more reliable estimate of the frequency of LSCs, however, is the co-localization of double-positive cells (p63+ BrdU+) (Fig. 4D, E). By this estimate, there was a significant reduction in LSCs following induction of a LSCD by either ethanol or rotating burr as compared to naïve mice. Importantly, there were significantly more double-positive (p63+ BrdU+) residual LSCs in LSCD mice with intact EBM compared to those without EBM (approximately 7.5-fold increase (*p* < 0.05) (Fig. 4D, E). These results indicate that total corneal de-epithelization performed using ethanol treatment results in less LSC exhaustion as compared to the rotating burr method.

### 3.4. Total corneal de-epithelization with or without EBM resulted in a complete LSCD

To ensure that both methods of inducing a LSCD (ethanol treatment and rotating burr) induced a complete LSCD, the following experiment was performed. Loss of corneal epithelial cells and infiltration of conjunctival epithelial cells is one of the defining characteristics of LSCD (Ruan et al., 2021). Therefore, we used CK19 (conjunctival epithelium-specific marker) (Donisi et al., 2003) staining to detect infiltrating conjunctival epithelial cells and CK12 (cornea epithelium-specific marker) to detect remaining corneal epithelial cells (Tanifuji-Terai et al., 2006) in eyes collected 14 days after total corneal de-epithelization. As expected, after de-epithelialization by either rotating burr (EBM-) or ethanol treatment (EBM+), there were no detectable CK12+ cells in either the central cornea or limbus (Fig.5A-D), indicating both techniques produced a complete LSCD. In both the EBM- and EBM+ groups, the loss of CK12+ corneal epithelial cells coincided with the presence of CK19+ infiltrating conjunctival epithelial cells, however, there was a slight decrease in the frequency of CK19+ cells in the mice receiving the ethanol treatment (EBM+) (Fig.5A-D).

**Fig. 5.**
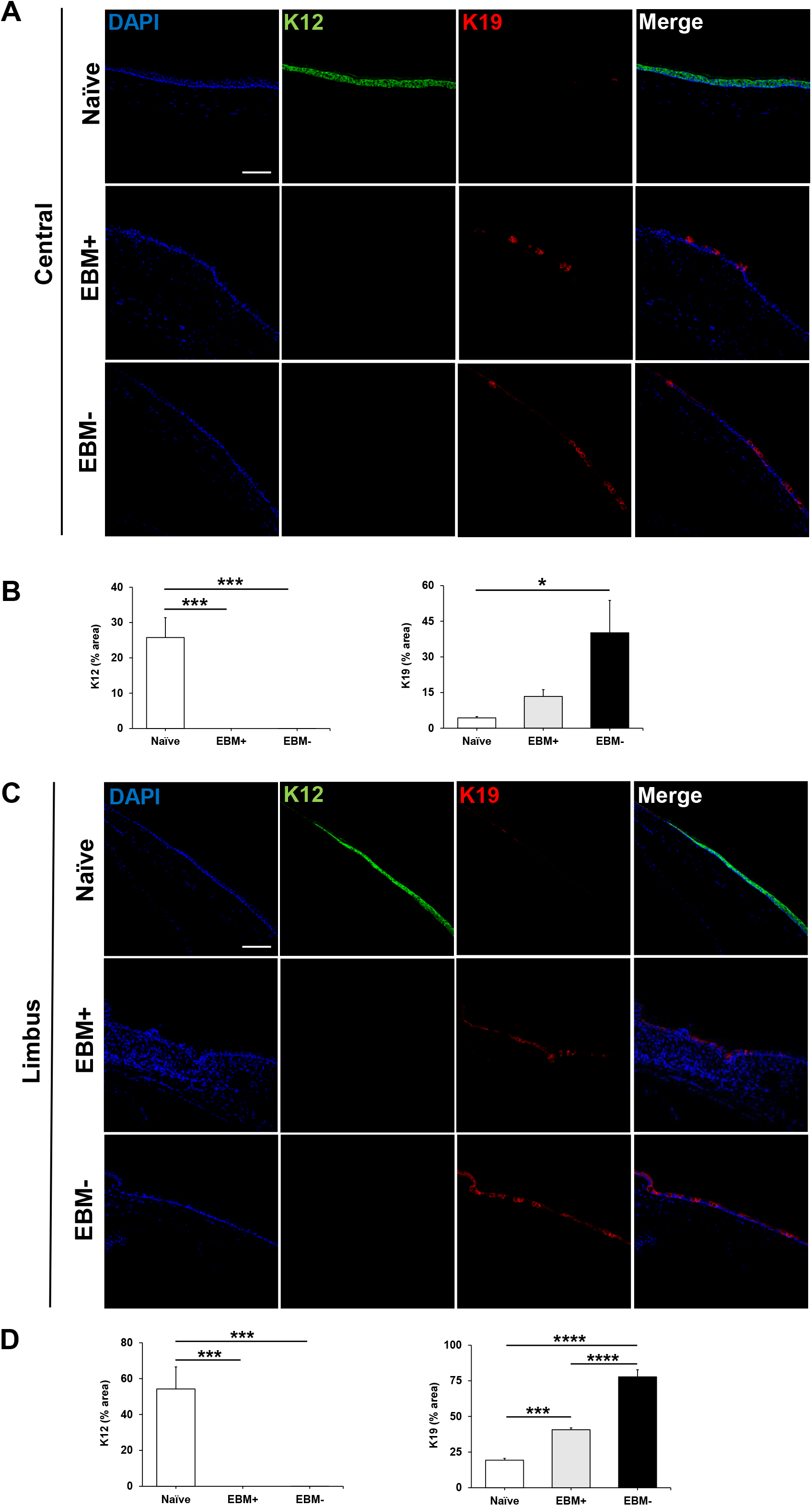
Total corneal de-epithelization with or without the EBM resulted in a complete LSCD. Representative CK12 and CK19 immunostaining image of the central cornea (A, B) and limbus (C, D) 14 days after total corneal de-epithelization in the EBM+ and EBM-groups. Quantitative graphs of the percentage of CK12 and CK19 stained cells in the cornea epithelium in the EBM+ and EBM-groups (n = 5 mice/group). Scale bars equal 100 μm. All data are presented as mean ± s.e.m. All statistical analyses were performed using a one-way analysis of variance followed by the Tukey-Kramer test. \**p* < 0.05; \*\**p* < 0.01; \*\*\**p* < 0.001; \*\*\*\**p* < 0.0001. EBM, epithelial basement membrane.CK12, cytokeratin 12. CK19, cytokeratin 19.

### 3.5. Total corneal de-epithelization with an intact EBM and residual LSCs coincides with reduced inflammation and corneal thickening

To evaluate corneal inflammation caused by total corneal de-epithelization, we measured CCT in EBM+ and EBM-groups every other day for two weeks. CCT was increased after de-epithelization in both EBM+ and EBM-groups compared to untreated normal controls. However, CCT was about two times greater (*p* < 0.05) in the EBM-group as compared with the EBM+ group on days 6-14 post-de-epithelization (Fig. 6A, B). Corneas were collected ten days after de-epithelization by rotating burr or ethanol treatment to quantitate pro-inflammatory factors (transforming growth factor-beta 1 (TGF-β1), interleukin-1 beta (IL-1β), and matrix metallopeptidase 9 (MMP-9)) (Wilson et al., 2020; AbuSamra et al., 2019). Naïve mice served as negative controls. De-epithelization by rotating burr that destroyed the EBM coincided with a significant increase in the mRNA levels of TGF-β1, IL-1β, and MMP-9, as compared with the negative controls (Fig. 6C). By contrast, de-epithelization by ethanol treatment that retained the EBM and possessed residual LSCs, showed significantly less mRNA for TGF-β1, IL-1β, and MMP-9, as compared with the EBM-group and no significant increase as compared to the negative controls (Fig. 6C).

**Fig. 6.**
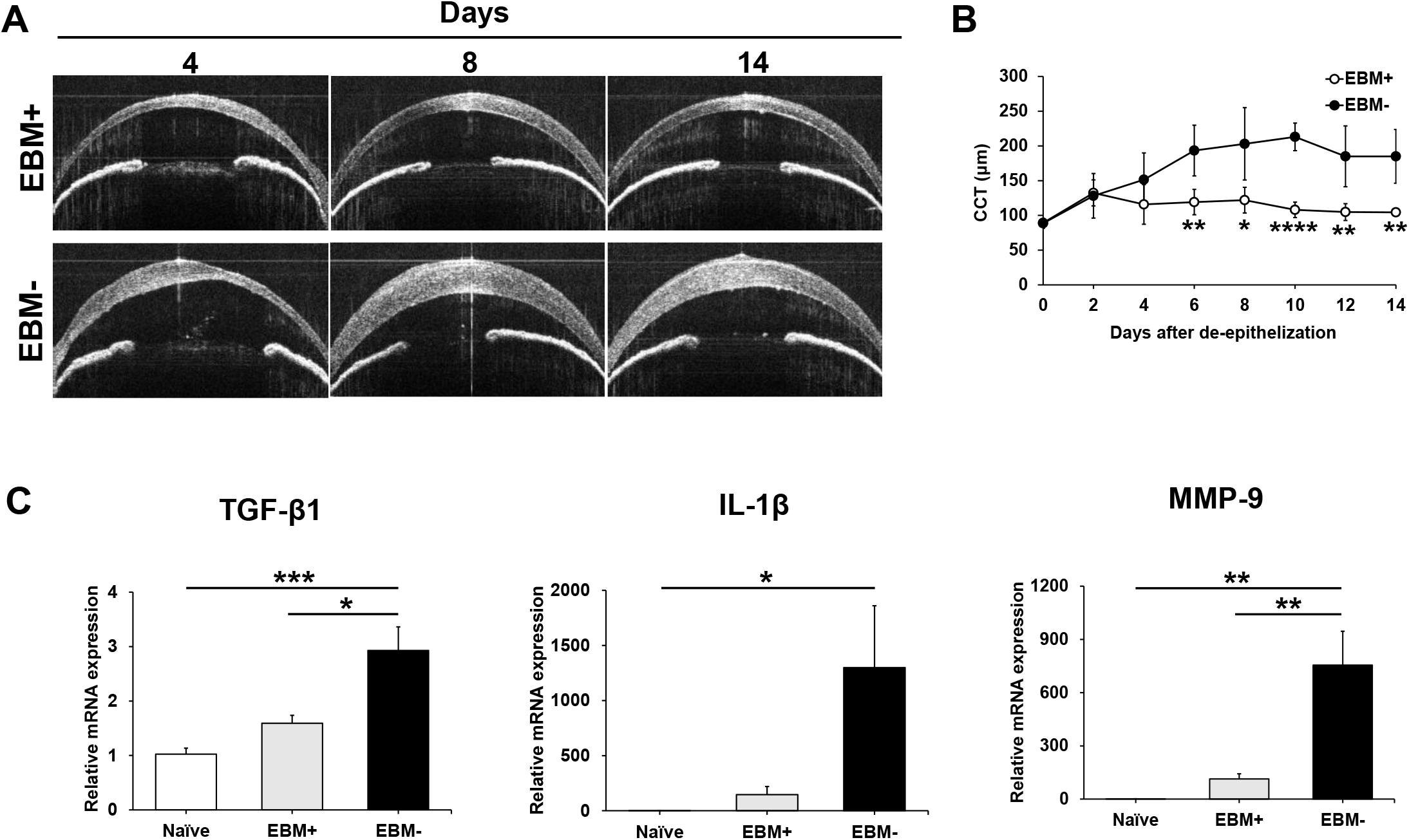
Total corneal de-epithelization with an intact EBM coincides with less pro-inflammatory factors and a reduced central cornea thickness. (A, B) Representative optical coherence tomography images on 4, 8, and 14 days after total corneal de-epithelization in the EBM+ and EBM-groups. Quantitative graph of the central corneal thickness (CCT) over time in the EBM+ and EBM-groups (n = 5 mice/group). Data are presented as mean ± S.D. Statistical analysis was performed using Student’s t-test. \**p* < 0.05; \*\**p* < 0.01; \*\*\*\**p* < 0.0001. (C) mRNA levels of TGF-β1, IL-1β, and MMP-9 in the cornea ten days after total corneal de-epithelization in the EBM+ and EBM-groups (n = 5 mice/group). Data are presented as mean ± s.e.m. Statistical analysis was performed using a one-way analysis of variance followed by the Tukey-Kramer test. \**p* < 0.05; \*\**p* < 0.01; \*\*\**p* < 0.001. EBM, epithelial basement membrane. TGF-β1, transforming growth factor beta 1. IL-1β, interleukin 1 beta. MMP-9; matrix metallopeptidase 9.

## 4. Discussion

We developed a LSCD model induced by performing total limbus-to-limbus corneal de-epithelization using an eye spear after loosening the adhesions between the epithelium and basement membrane with ethanol. In this model, we retained the EBM even after total corneal de-epithelization. The presence of an intact EBM significantly contributed to promoting wound closure, reduced levels of pro-inflammatory cytokines (IL-1β, TGFβ1, and MMP-9), and retained a small but significant number of residual LSCs as measured by the number of double-positive p63+ BrdU+ cells. In this LSCD model, we placed a cylindrical filter paper (inner and outer diameter: 3 and 4 mm) outside the limbus. This enabled us to prevent ethanol spillover to the outside of the limbus because the cornea diameter of 6-8 weeks old C57BL/6 mice is 2.6 mm (Henriksson et al., 2009). Furthermore, loosening the binding between basal epithelial cells and the EBM using ethanol allows us to scrape off the epithelium more efficiently and uniformly without disturbing the EBM. All these innovations led to making a highly reproducible LSCD model with a small number of residual LSCs.

After total corneal de-epithelization, the EBM+ group had slightly less corneal opacity than the EBM-group (Fig. 3C, D). A previous study showed that disrupting the EBM allowed epithelial-derived TGFβ1 to reach the stroma, resulting in the differentiation of stromal fibroblast precursor cells to differentiated fibroblasts. This phenomenon was considered a significant cause of corneal opacity (Singh V et al., 2014). Considering this, in the EBM+ group, inhibiting TGFβ1 from reaching the stroma might contribute to the slight reduction in corneal opacity compared to the EBM-group.

Significantly less corneal edema was observed in the EBM+ group after de-epithelization than in the EBM-group (Fig. 6A, B). This coincided with reduced inflammation as measured by decreased levels of IL-1β, TGFβ1, and MMP-9 mRNA in the cornea (Fig. 6C). EBM disruption enables the cornea epithelium to contact stromal fibroblasts directly, and a previous study showed this direct interaction between epithelial cells and stromal fibroblasts induced IL-1β secretion by fibroblasts (AbuSamra et al., 2019). Considering this finding, in the EBM+ group, blocking the interaction between epithelium and stroma might bring about a decrease in the production of IL-1β mRNA, resulting in less corneal inflammation. Since inflammation in the LSC microenvironment is considered to be one of the major causes of LSC exhaustion (Ruan et al., 2021), reduced inflammation may contribute to residual LSCs in complete LSCD mice induced using ethanol.

In summary, we made a highly reproducible LSCD murine model with residual LSCs by performing total corneal de-epithelization using ethanol. This LSCD murine model could be a useful animal model for determining the function of residual LSCs and whether they can be rejuvenated and expanded to reestablish and maintain the corneal epithelium.

## Acknowledgments

We thank the technical staff at the Schepens Eye Research Institute of Massachusetts Eye and Ear, Harvard Medical School, for supporting this research. This study was supported by the National Institute of Health/National Eye Institute: Grant R01EY025794 (Markus H. Frank, Natasha Y. Frank, and Bruce R. Ksander) and Core Grant P30EY003790 (Patricia A. D’amore).

## SUPPLEMENTARY DATA

**Fig. S1.**
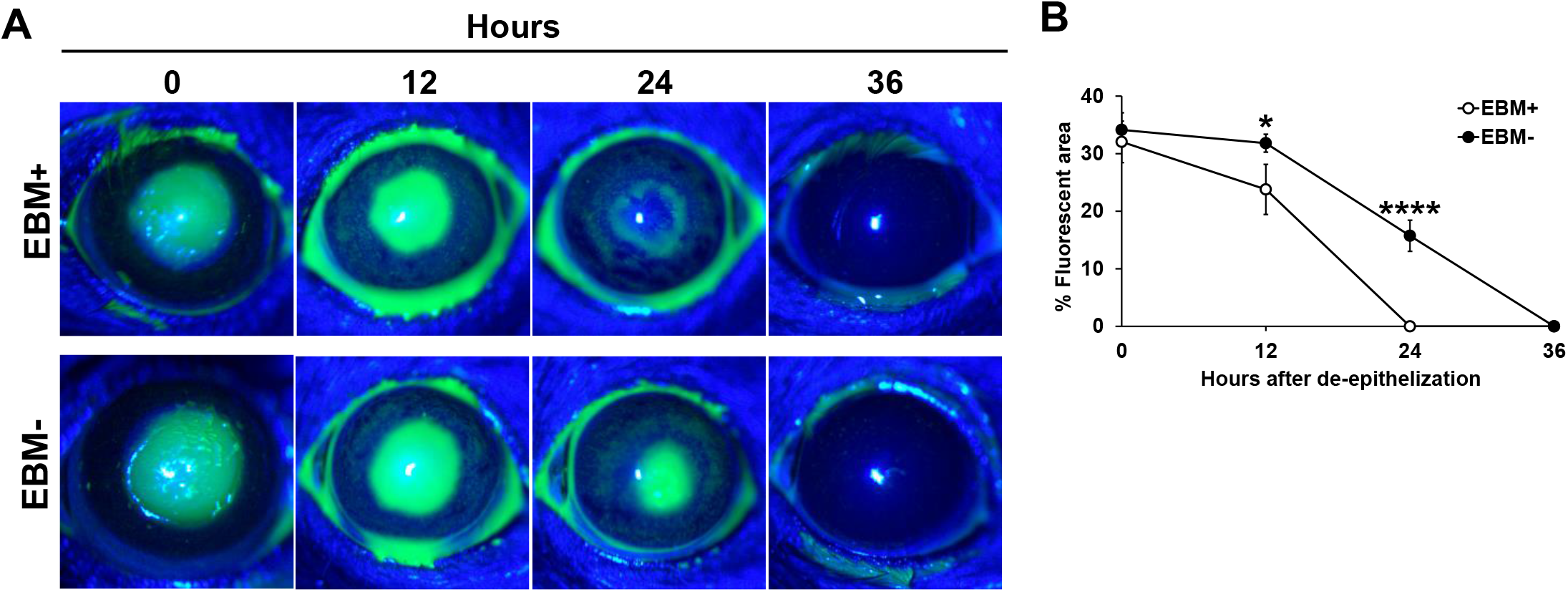
The presence of EBM after central corneal de-epithelization is critical for promoting cornea epithelial wound healing. (A, B) Representative fluorescein-stained cornea photographs on 12, 24, and 36 hours after central corneal de-epithelization in the EBM+ and EBM-groups. Quantitative graph of the % fluorescein-stained area over time in the EBM+ and EBM-groups (n = 4 mice/group). Data are presented as mean ± S.D. Statistical analysis was performed using Student’s t-test. \**p* < 0.05; \*\*\*\**p* < 0.0001. EBM, epithelial basement membrane.

**Table. S1. The primer sequence used in this study**.

## References

AbuSamra, D. B., Mauris, J., & Argüeso, P. (2019). Galectin-3 initiates epithelial-stromal paracrine signaling to shape the proteolytic microenvironment during corneal repair. Science signaling, 12, eaaw7095.

Afsharkhamseh, N., Movahedan, A., Gidfar, S., Huvard, M., Wasielewski, L., Milani, B. Y., Eslani, M., & Djalilian, A. R. (2016). Stability of limbal stem cell deficiency after mechanical and thermal injuries in mice. Experimental eye research, 145, 88–92.

Browning, A. C., Shah, S., Dua, H. S., Maharajan, S. V., Gray, T., & Bragheeth, M. A. (2003). Alcohol debridement of the corneal epithelium in PRK and LASEK: an electron microscopic study. Investigative ophthalmology & visual science, 44, 510–513.

Chan, Eric, Qihua Le, Andres Codriansky, Jiaxu Hong, Jianjian Xu, and Sophie X. Deng. (2016). Existence of Normal Limbal Epithelium in Eyes with Clinical Signs of Total Limbal Stem Cell Deficiency. Cornea, 35, 1483–87.

Delic, N. C., Cai, J. R., Watson, S. L., Downie, L. E., & Di Girolamo, N. (2022). Evaluating the clinical translational relevance of animal models for limbal stem cell deficiency: A systematic review. The ocular surface, 23, 169–183.

Deng, Sophie X., Vincent Borderie, Clara C. Chan, Reza Dana, Francisco C. Figueiredo, José A. P. Gomes, Graziella Pellegrini, Shigeto Shimmura, Friedrich E. Kruse, and The International Limbal Stem Cell Deficiency Working Group. (2019). Global Consensus on Definition, Classification, Diagnosis, and Staging of Limbal Stem Cell Deficiency. Cornea, 38, 364–75.

Deng, Sophie X., Friedrich Kruse, José A. P. Gomes, Clara C. Chan, Sheraz Daya, Reza Dana, Francisco C. Figueiredo, et al. (2020). Global Consensus on the Management of Limbal Stem Cell Deficiency. Cornea, 39, 1291–1302.

Donisi, P. M., Rama, P., Fasolo, A., & Ponzin, D. (2003). Analysis of limbal stem cell deficiency by corneal impression cytology. Cornea, 22, 533–538.

Dua, H. S., Deshmukh, R., Ting, D. S. J., Wilde, C., Nubile, M., Mastropasqua, L., & Said, D. G. (2021). Topical use of alcohol in ophthalmology - Diagnostic and therapeutic indications. The ocular surface, 21, 1–15.

Espana, E. M., Grueterich, M., Mateo, A., Romano, A. C., Yee, S. B., Yee, R. W., & Tseng, S. C. (2003). Cleavage of corneal basement membrane components by ethanol exposure in laser-assisted subepithelial keratectomy. Journal of cataract and refractive surgery, 29, 1192–1197.

Henriksson, J. T., McDermott, A. M., & Bergmanson, J. P. (2009). Dimensions and morphology of the cornea in three strains of mice. Investigative ophthalmology & visual science, 50, 3648–3654.

Kate, A., & Basu, S. (2022). A Review of the Diagnosis and Treatment of Limbal Stem Cell Deficiency. Frontiers in medicine, 9, 836009.

Ksander, B. R., Kolovou, P. E., Wilson, B. J., Saab, K. R., Guo, Q., Ma, J., McGuire, S. P., Gregory, M. S., Vincent, W. J., Perez, V. L., Cruz-Guilloty, F., Kao, W. W., Call, M. K., Tucker, B. A., Zhan, Q., Murphy, G. F., Lathrop, K. L., Alt, C., Mortensen, L. J., Lin, C. P., … Frank, N. Y. (2014). ABCB5 is a limbal stem cell gene required for corneal development and repair. Nature, 511, 353–357.

Lasagni Vitar, R., Triani, F., Barbariga, M., Fonteyne, P., Rama, P., & Ferrari, G. (2022). Substance P/neurokinin-1 receptor pathway blockade ameliorates limbal stem cell deficiency by modulating mTOR pathway and preventing cell senescence. Stem cell reports, 17, 849–863.

Li, F. J., Nili, E., Lau, C., Richardson, N. A., Walshe, J., Barnett, N. L., Cronin, B. G., Hirst, L. W., Schwab, I. R., Chirila, T. V., & Harkin, D. G. (2016). Evaluation of the AlgerBrush II rotating burr as a tool for inducing ocular surface failure in the New Zealand White rabbit. Experimental eye research, 147, 1–11.

Mahmood, N., Suh, T. C., Ali, K. M., Sefat, E., Jahan, U. M., Huang, Y., Gilger, B. C., & Gluck, J. M. (2022). Induced Pluripotent Stem Cell-Derived Corneal Cells: Current Status and Application. Stem cell reviews and reports, 18, 2817–2832.

Pellegrini, G., Dellambra, E., Golisano, O., Martinelli, E., Fantozzi, I., Bondanza, S., Ponzin, D., McKeon, F., & De Luca, M. (2001). p63 identifies keratinocyte stem cells. Proceedings of the National Academy of Sciences of the United States of America, 98, 3156–3161.

Ruan, Y., Jiang, S., Musayeva, A., Pfeiffer, N., & Gericke, A. (2021). Corneal Epithelial Stem Cells-Physiology, Pathophysiology and Therapeutic Options. Cells, 10, 2302.

Sartaj, R., Zhang, C., Wan, P., Pasha, Z., Guaiquil, V., Liu, A., Liu, J., Luo, Y., Fuchs, E., & Rosenblatt, M. I. (2017). Characterization of slow cycling corneal limbal epithelial cells identifies putative stem cell markers. Scientific reports, 7, 3793.

Satake, Yoshiyuki, Murat Dogru, Takefumi Yamaguchi, Daisuke Tomida, Masatoshi Hirayama, and Jun Shimazaki. (2013). Immunological Rejection Following Allogeneic Cultivated Limbal Epithelial Transplantation. JAMA Ophthalmology, 131, 920–22.

Singh, V., Jaini, R., Torricelli, A. A., Santhiago, M. R., Singh, N., Ambati, B. K., & Wilson, S. E. (2014). TGFβ and PDGF-B signaling blockade inhibits myofibroblast development from both bone marrow-derived and keratocyte-derived precursor cells in vivo. Experimental eye research, 121, 35–40.

Sonoda, Y., & Streilein, J. W. (1992). Orthotopic corneal transplantation in mice--evidence that the immunogenetic rules of rejection do not apply. Transplantation, 54, 694–704.

Tanifuji-Terai, N., Terai, K., Hayashi, Y., Chikama, T., & Kao, W. W. (2006). Expression of keratin 12 and maturation of corneal epithelium during development and postnatal growth. Investigative ophthalmology & visual science, 47, 545–551.

Wilson, S. E., Torricelli, A. A. M., & Marino, G. K. (2020). Corneal epithelial basement membrane: Structure, function and regeneration. Experimental eye research, 194, 108002.

